# Diet diversification shapes broad-scale distribution patterns in European bats

**DOI:** 10.1101/704759

**Authors:** Antton Alberdi, Orly Razgour, Ostaizka Aizpurua, Roberto Novella-Fernandez, Joxerra Aihartza, Ivana Budinski, Inazio Garin, Carlos Ibáñez, Eñaut Izagirre, Hugo Rebelo, Danilo Russo, Anton Vlaschenko, Violeta Zhelyazkova, Vida Zrncic, M Thomas P Gilbert

## Abstract

Large-scale species’ distributions have been traditionally attributed to physiological traits related to abiotic factors, while behavioural features linked to biotic interactions have received little attention. We tested the relationship between trophic and spatial niche breadths through combining species distribution modelling with dietary DNA metabarcoding of over 400 bats sampled across Europe belonging to seven species. Our results point to a causality cascade between hunting plasticity, trophic niche breadth and spatial niche breadth, and thus indicate that behavioral plasticity and dietary diversification can contribute to shaping broad-scale species distributions.

## Introduction

The characterisation and comparison of trophic niches have been core topics in animal ecology since Hutchinson’s conceptualisation of the ecological niche (1). While traditionally considered important only for local-scale species interactions (2), dietary features have also proven relevant for broader scale species distributions (3). Nevertheless, the link between species’ trophic niches and large-scale distribution patterns is still inconclusive (4). This could be because diet analyses have traditionally relied on particular diversity metrics that overlook some components of dietary diversity, and because until recently, methodological constraints have limited the possibility of performing broad-scale high-resolution diet studies (5).

Following the advent of high throughput DNA sequencing-based tools, it is now possible to characterise dietary niches across much larger sample sizes, and at levels of detail never seen before (6). DNA-based diversity assessment also enables comprehensive analysis of trophic variation through considering different components of dietary diversity, such as richness (how many prey are consumed), evenness (the balance of the relative consumption of each prey) and regularity (the degree of similarity across consumed prey) (7,8). Richness, evenness and regularity metrics are positively associated with performance in several ecological systems (9). Hence, we hypothesised that these metrics applied to trophic niches also impact the capacity of animals to thrive in a wider range of environmental conditions, and hence trophic niche breadth could potentially contribute to the shaping of species’ distributions.

To test this, we contrasted broad-scale dietary and spatial niches of a vertebrate system, namely the European bat community. Bats provide an excellent opportunity for understanding dietary and spatial diversity patterns due to their spatial variability and well-studied behavioural traits (10). We collected faecal samples from over 400 bats representing seven species captured at 40 locations scattered across the European continent. Faeces of each individual bat were independently analysed through DNA metabarcoding and high throughput sequencing by using two complementary primer sets and three replicates per primer. We used the statistical framework recently developed around Hill numbers (11) to contrast trophic niche measures based on richness (dR), richness+evenness (dRE) and richness+evenness+regularity (dRER). The different species-level trophic niche measures were then statistically related to spatial niche breadth metrics as measured by species distribution modelling, as well as to a range of behavioural traits to assess the causal directionality between dietary diversification and spatial niche expansion.

## Results

### The trophic niche of European bats is dominated by Lepidoptera and Diptera

After applying all quality filters, the dataset included dietary information of 355 individual bats belonging to seven species (DNA sequencing details in Table S4). Using two primer sets, we detected over 3000 different prey taxa belonging to 29 arthropod orders (Fig. 1A), though the pandiet of European bats was dominated by Lepidoptera and Diptera (Fig. 1B). Our results complement the existing broad-scale molecular dietary data of *Miniopterus schreibersii* (12), and provide the first geographically widespread molecular insights into the dietary ecology of *Myotis daubentonii, M. myotis*, *M. emarginatus*, *M. capaccinii*, *Rhinolophus euryale* and *R. ferrumequinum*, which had only been studied at local scales previously (13–15).

**Figure 1.**
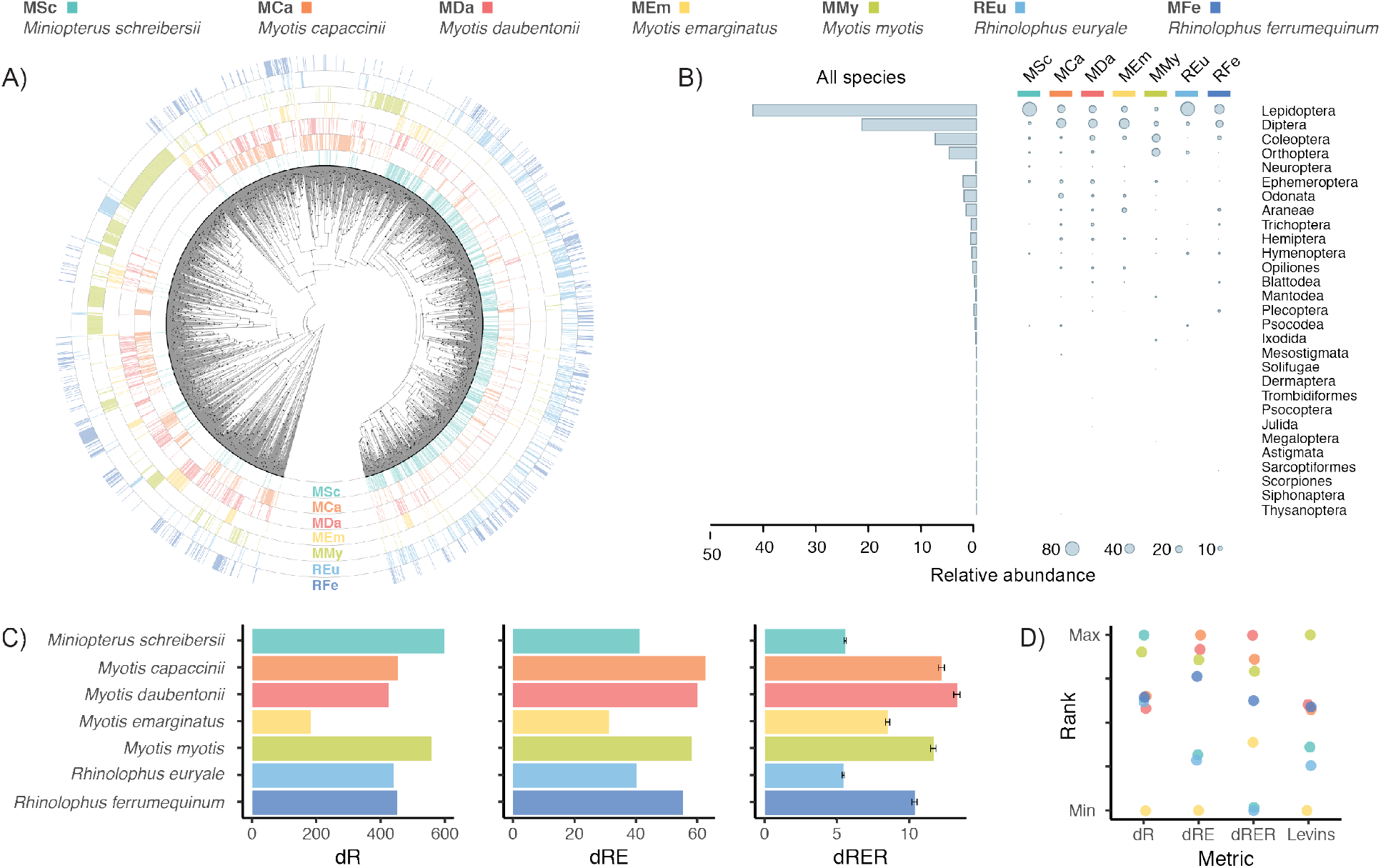
Dietary diversity statistics of the analysed bat species. (A) Radial phylogenetic tree of prey detected using the Zeale primers and their occurrence patterns in each of the studied bats. A higher resolution image (Fig. S1) and the homologous figure built from the Epp data (Fig. S2) are available in the Supplementary Information. (B) Overall and predator species-specific representation of the arthropod taxonomic orders. (C) Dietary niche breadth measures accounting for richness (dR), richness+evenness (dRE) and richness+evenness+regularity (dRER). The error bars (±SE) of dRER indicate the dispersion of the trophic niche breadths yielded when using different prey phylogenetic trees (N=50) sampled from the Bayesian MCMC. (D) Species ranks according to the dietary niche breadth measure type. Levin’s index is also included for being the most common metric employed in the literature.

### Trophic niche differences depend on the components of diversity accounted for

Trophic niche breadth measures (Fig. 1C), and the species ranks derived from them (Fig. 1D), were different depending on the components of diversity considered. Similar contrasting results have also been reported in other systems (16), due to the fact that each diversity component might be driven by different ecological forces (17). For instance, the trophic niches of *M. schreibersii* and *R. euryale* showed similar dRE values, yet the contribution of richness and relative evenness components differed. The dietary richness of *M. schreibersii* was almost 40% larger than *R. euryale*’s, while the evenness factor of *R. euryale* was almost 30% higher than that of *M. schreibersii.* These differences could be explained by i) the larger home range of *M. schreibersii* compared to *R. euryale* (18,19), which might expose the former to more prey species —thus increasing dietary richness, and ii) the higher incidence of a few locally abundant pest moth species in the diet of *M. schreibersii* than in that of *R. euryale* (12,14) —yielding lower relative evenness. As reported for other systems (20), trophic niche differences would be overlooked if the niche breadth analyses were limited to a single diversity metric.

### Trophic niche breadth explains spatial niche breadth

The distribution models generated (Table S5) to test whether trophic niche breadth correlates with spatial niche breadth yielded different spatial projections (Fig. S3) and niche breadth measures for each species. We found that the trophic niche breadth measures accounting for all diversity components (dRER) were positively correlated with the two spatial niche breadth metrics computed, both for each primer-specific dataset (Table S8, Fig. S4) and the overall averaged dataset (Levins’ B1: Pearson’s r = 0.85; t = 29.57, df = 348, p-value < 0.001; Levins’ B2 (Fig. 2A): Pearson’s r = 0.80; t = 24.58, df = 348, p-value < 0.001). The species that consume a wider variety of distinct prey are the ones that exhibit broader spatial niches.

**Figure 2.**
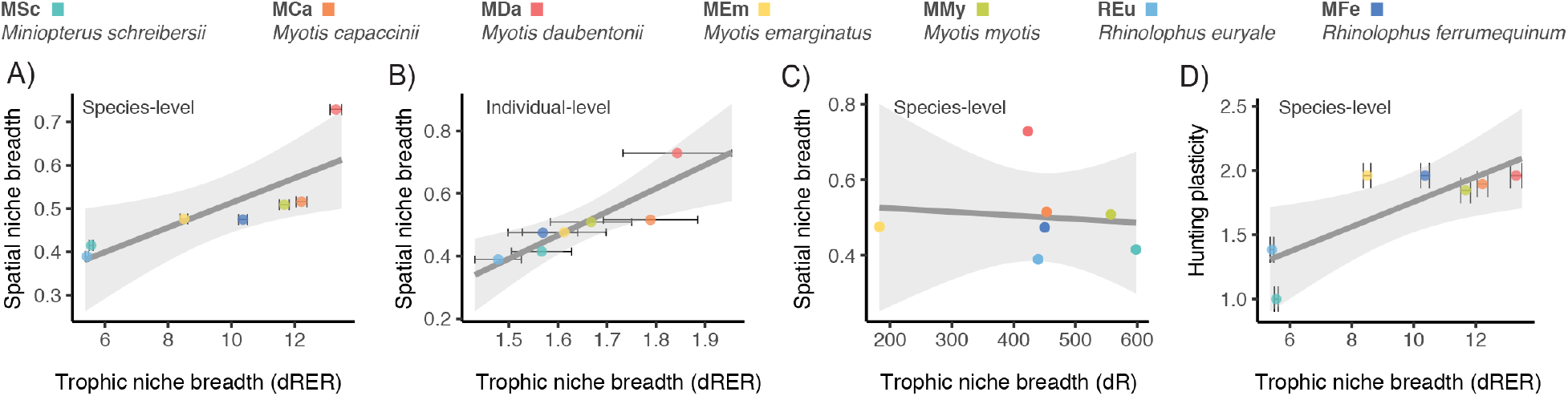
Relation of species trophic niche breadth measures with spatial niche breadth and hunting plasticity. (A) Species-level relation between niche breadth measures accounting for the three components of diversity (dRER: richness+evenness+regularity) and spatial niche breadths of the respective species. (B) Individual-level relation between dRER niche breadth measures and spatial niche breadths of the respective species. (C) Species-level relation between niche breadth measures accounting for only one component of diversity (dR: richness) and spatial niche breadths of the respective species. (D) Species-level relation between dRER niche breadth measures and hunting plasticity of the respective species. Dots indicate mean values per bat species. Note that error bars (±SE) at the species-level charts indicate the dispersion of the different trophic niche breadth values yielded from the 50 iterations run with different prey phylogenetic trees to account for phylogenetic uncertainty. In contrast, the error bars at the individual-level chart indicates the dispersion of the individual bats’ trophic niche breadth values. Chart C does not contain error bars as for dR a single niche breadth measure was computed for each species.

We performed a range of analyses to further assess the plausibility of the causal relationship between dietary diversification and spatial niche breadth. We first assessed whether dietary breadths could be broadened passively as a result of spatial niche expansion (21). If the ability to thrive in more distinct environments was driving dietary expansion, we would expect species dietary breadth to be driven by beta diversity, i.e. dietary differences across individuals within species. However, we observed that the correlation between trophic and spatial niche breadth remained significant (Pearson’s r = 0.20,t = 3.78, df = 353, p-value < 0.001; Fig. 2B), which highlights the relevance of alpha (individual) dietary diversity. Additionally, if dietary breadths were passively broadened due to spatial niche expansion, we would expect species dietary richness to also increase, as predators are exposed to a larger variety of potential prey. Nevertheless, we observed that the correlation between trophic and spatial niche breadths disappeared when relative evenness and regularity components were removed (Levins’ B1: Pearson’s r = −0.09; t = −0.20, df = 5, p-value = 0.846; Fig. 2C). Consequently, these two observations rule out the possibility that dietary breadths are passively broadened as a result of spatial niche expansion.

We then investigated the relation between different behavioural traits (hunting plasticity, habitat use diversity and roosting plasticity) and spatial niche breadths to assess whether a third factor could be shaping both dietary and spatial niches. We found no significant correlation between any of the analysed traits and spatial niche breadth (Table S9), yet we found that hunting plasticity is positively related with the trophic niche breadth of predator species (Pearson’s r = 0.79, t = 24.347, df = 348, p-value < 0.001; Fig. 2D). This suggests that the ability to use a more diverse range of hunting strategies, such as capturing prey from the ground, foliage or water surface, in addition to hunting flying prey (10), broadens the functional spectrum of captured prey, which in turn widens the spatial niche.

## Discussion

This is the first time that high-resolution trophic (DNA metabarcoding) and spatial (Species Distribution Modelling) niche characterisation identify diet as a driving factor of broad-scale spatial patterns. While ecological niche breadth has been previously shown to predict geographical range sizes, trophic niche breadth has so far only been associated with broad-scale spatial patterns in arthropods (4). The causality cascade between hunting plasticity, dietary niche breadth and spatial niche breadth seems plausible and is ecologically meaningful. Hunting plasticity has previously been linked to adaptability (22), and could directly affect the fitness of bats, for instance by enabling shifting diets when specific prey types become scarce. Trophic niche breadth could also indirectly affect the fitness of bats by, for example, fostering gut microbiome diversification and dynamism, which have been associated with adaptation capacity in vertebrates (23).

Our results contradict the Eltonian noise hypothesis, which proposes that biotic interactions do not affect species distributions at large geographical scales (2). However, in this case spatial patterns would not primarily depend on the availability of resources as previously shown (3), but on the inherent behavioural properties of predators. It is noteworthy that all the patterns we found in this study were recovered from trophic niche measures that considered all three components of diversity. No diversity component alone was correlated with either hunting plasticity or spatial niche breadth, which highlights the importance of accounting for relative evenness and regularity of prey when measuring trophic niches. It is also remarkable that a single snapshot of the diet of individual bats was enough to recover a clear link between trophic and spatial niche breadth, as niche patterns are not always coupled at individual and population levels (24). Overall, our study demonstrates the potential of combining environmental DNA with species distribution modelling and behavioural ecology, to unveil broad-scale ecological patterns and links between different components of the ecological niche of species. Finally, our results also highlight the relevance of diet in shaping broad-scale animal distributions, which supports the use of behavioural plasticity as a relevant feature to predict species’ range shifts in response to climate change (25).

## Methods

### Data collection and generation

We collected droppings from 402 individual bats captured in 40 locations distributed across Europe (Table S1), in June-October of 2015-2017. The droppings belonged to seven species: *Miniopterus schreibersii* (MSc), *Myotis capaccinii* (MCa), *Myotis daubentonii* (MDa), *Myotis emarginatus* (MEm), *Myotis myotis* (MMy), *Rhinolophus euryale* (REu) and *Rhinolophus ferrumequinum* (RFe). Using a randomised setup, DNA was extracted from all individual samples and amplified in three replicates using two primer pairs, referred to as Zeale (26) and Epp (27). Amplicons were purified, pooled and built into libraries before Illumina MiSeq sequencing. To ensure maximum DNA sequence reliability, only high quality sequences that appeared in at least two of the three PCR replicates were retained, and sequences identical to those detected in the extraction and library blanks of the corresponding processing batch of each sample were removed. Appropriate sampling depth per sample was ensured by discarding samples with insufficient sequencing depth as assessed by rarefaction curves and curvature indexes. DNA sequences were clustered into operational taxonomic units (OTUs) based on 98% identity following Alberdi et al. (28) and taxonomy was assigned by aligning the OTU representative sequences to the Genbank nt (29) —and in the case of Zeale also BOLD (30)— databases. Full details of the field, laboratory and bioinformatics methodologies are reported in the Supplementary Information and Supplementary Code 1.

### Data analysis

Diversity analyses were carried out using the R package hilldiv (31) based on abundance-based Hill numbers (32,33). The Hill numbers framework enables i) the relative weight given to abundant and rare OTUs to be modulated through a single parameter, namely the order of diversity q (32), and ii) the similarity level across OTUs to be overlooked or accounted for when computing diversity. Although functional diversities can be computed using Hill numbers (34), given the infeasibility of gathering ecological trait information of thousands of prey items, OTU phylogenies were employed as proxies of ecological resemblance across OTU. Hence, dR (richness) was computed as the neutral Hill number of order of diversity q=0; dRE (richness+evenness) was computed as the neutral Hill number of order of diversity q=1 —i.e. Shannon diversity— and dRER (richness+evenness+regularity) was computed as the phylogenetic Hill number of order of diversity q=1. Phylogenetic Hill numbers were computed based on Bayesian phylogenies generated from metabarcoding DNA sequences, and the analyses accounted for the phylogenetic uncertainty of generated trees, as detailed in Supplementary Information. Sample size appropriateness was assessed by comparing observed vs. estimated trophic niche breadth values (Table S14).

Predator species’ hunting strategy, habitat-use and roosting data were gathered from 45 articles available in the literature (Tables S12-14). Plasticity indices were computed by means of Shannon diversity of ecological traits. The species distributions models that characterised the spatial niche of predator species were generated using BIOMOD (35), and niche breadths were measured by means of Levins’ B1 and B2 metrics (36) based on the spatial projections using the R version ENMTools (37). For all statistical tests, significance threshold was set at p=0.05. All statistical analyses were performed in R (38) after averaging the results yielded by both primers unless otherwise stated.

## Data availability

The datasets generated during and/or analysed during the current study are available in the Dryad repository (ref. [to be included in the reference list when a DOI is available]).

## Code availability

The bash, python and R scripts used for analysing the data during the current study are available in the Supplementary Files as Supplementary Code 1 (DNA metabarcoding), Supplementary Code 2 (Species Distribution Modelling) and Supplementary Code 3 (ecological niche statistical analyses).

## Author contributions

A.A., O.A. and M.T.P.G designed the study. All authors participated in the sample and data collection. A.A. and O.A. performed the laboratory procedures. O.R. and R.N.F. carried out the species distribution modelling. O.A. performed the ecological trait analyses. A.A. performed the DNA metabarcoding and statistical analyses. A.A. wrote the manuscript. All authors contributed to and approved the final version.

## Acknowledgements

A.A. was supported by Lundbeckfonden (R250-2017-1351) and the Danish Council for Independent Research (DFF 5051-00033). O.R. was supported by a NERC Independent Research Fellowship (NE/M018660/1), and O.A. was supported by the Carlsberg Foundation’s Postdoctoral Fellowship (CF15-0619). M.T.P.G. acknowledges ERC Consolidator Grant (681396-Extinction Genomics). We are grateful to Fiona Mathews, Daniel Whitby, Roger Ransome, Matt Cook and Martina Spada for providing samples; and Aitor Arrizabalaga, Lide Jimenez, Vilalii Hukov, Olena Holovchenko, Vanessa Mata, and Branka Pejić for assistance in the field work.

