## Supplementary Information for "Diet diversification shapes broad-scale distribution patterns in European bats"

#### Table of contents

**Methods** **2**
- 1. Sampling and sample storage 2
- 2. Laboratory procedures 2
  - 2.1 DNA extraction 2
  - 2.2 Quantitative PCR (qPCR) screening 3
  - 2.3 PCR amplification 4
  - 2.4 Amplicon visualization, pooling and purification 4
  - 2.5 Library preparation and sequencing 5
- 3. Metabarcoding bioinformatic procedures 5
- 4. Species distribution modelling 6
- 5. Statistical analyses and visualisations 6

**Tables** **8**
- Table S1. Sampling sites. 8
- Table S2. PCR reaction recipes. 9
- Table S3. Thermal cycler programs 9
- Table S4. DNA sequence processing statistics 10
- Table S5. Species distribution model statistics. 10
- Table S6. Description of the variables used in the species distribution models and their sources. 11
- Table S7. List of publications used to extract species occurrence records. 12
- Table S8. Primer-specific relation between trophic (dRER) and spatial niche breadth 13
- Table S9. Relation between behavioral traits and spatial niche breadth 14
- Table S10. Relation between behavioral traits and dietary niche breadth 14
- Table S11. Correlation between observed and estimated diversities 14
- Table S12. Hunting strategy traits 15
- Table S13. Habitat use traits 16
- Table S14. Roosting traits 17

**Figures** **18**
- Figure S1. Trophic spectrum of the analysed species as retrieved from the Zeale primers 18
- Figure S2. Trophic spectrum of the analysed species as retrieved from the Epp primers 19
- Figure S3. Spatial projections of the niche models 20
- Figure S4. Relation of species trophic niche breadth measures with spatial niche breadth 21

**References** **22**

### Methods

#### 1. Sampling and sample storage

We collected droppings from individual bats in 40 locations distributed across Europe in summer 2015-2017. The mean geographical distance between sampling locations was 1398 km, with a maximum distance of 3707 km between Portugal and Ukraine (Table S1). We captured the bats in roost entrances using harp-traps and/or mist-nets when returning from foraging (1am-7am), which ensured rapid defecation. To avoid sample cross-contamination, each bat was kept separately in a clean, single-use, UV-radiation sterilised cotton bag for 15-20 minutes, then identified, sexed and aged before releasing it into the cave. Faecal pellets were collected from the bags and stored in 1.5 ml tubes filled with silica gel granules (Chameleon® C 1-3 mm, VWR) or absolute ethanol. Samples were kept dried and refrigerated (4-8 ºC) until they were transported to the laboratory, after which they were stored at -20 ºC. All captures were authorized according to the laws of the countries where they were carried out.

#### 2. Laboratory procedures

##### 2.1 DNA extraction

From each of the 402 individual bats, DNA was extracted from 1-3 bat droppings (approximate total weight: 20-30 mg) using the PowerSoil ® DNA Isolation Kit (MoBio, CA, USA) principally following the manufacturer's protocol (2016 version), but with some modifications. Each extraction round included 23 bat dropping samples and one negative extraction control. Extractions were performed in a dedicated pre-PCR laboratory. The exact employed protocol is detailed below:

1. From each individual bat, 1-5 droppings (20-30 mg) were added to the PowerBead Tubes containing silica beads and beads solution.
2. 60 µl of C1 solution were added to the tubes briefly vortexed.
3. The tubes were heated for 15 minutes at 65 ºC in an incubator.
4. Samples were bead-beaten for 10 minutes at frequency 1/20 on a Qiagen TissueLyser II.
5. Tubes were centrifuged at 13,000g for 3 minutes.
6. The supernatant (400-500 µl) were transferred to new 1.5 ml tubes.
7. 250 µl of C2 solution were added to the tubes followed by brief vortexing.

8. Tubes were incubated for 10 minutes at 4 ºC.
9. Tubes were centrifuged at 13,000g for 1 minute.
10. The supernatants (up to 600 µl) were transferred to new 1.5 ml tubes.
11. 200 µl of C3 solution were added followed by brief vortexing.
12. Tubes were incubated for 10 minutes at 4 ºC.
13. Tubes were centrifuged at 13,000g for 1 minute.
14. Supernatants (up to 750 µl) were transferred to new 1.5 ml tubes.
15. 1200 µl of C4 solution were added followed by brief vortexing.
16. 650 µl of the mixes were loaded onto spin filters and centrifuged at 8,000g for 1 minute.
17. Flowthroughs were discarded and the operations 16 and 17 repeated until loading all the mix.
18. 500 µl of C5 solution were added and centrifuged at 13,000g for 1 minute.
19. Flowthroughs were discarded and columns centrifuged again at 13,000g for 1 minute.
20. Spin filters were placed in new 2 ml collection tubes and add 50 µl of EB buffer were added to the filters.
21. Tubes were incubated for 15 minutes at 37-40 ºC.
22. Tubes were centrifuged at 13,000g for 1 minute and DNA extracts transferred to 1.5 ml low-bind tubes.

#### 2.2 Quantitative PCR (qPCR) screening

Before conductin tagged PCRs, for each of the two primer pairs employed, quantitative PCR (qPCR) screening with multiple DNA template volumes and dilutions was carried out in a subset of samples to (i) assess contamination of extraction blanks, (ii) determine the optimal cycle number for the subsequent PCRs, and (iii) estimate the maximum template amount for the following tagged PCR amplifications in which PCR inhibitory substances, copurified with the DNA, would not distort the amplification (1–3) . Initial SYBR green chemistry qPCR screenings were carried out on all extraction blanks and on a dilution series (3 µl, 2 µl, 1 µl, and 1:1, 1:5 and 1:10) of 28 bat faecal extracts (four per species) for each of the two primer sets. The most appropriate primer concentration for each reaction was also assessed by qPCR.

Quantitative PCRs were carried out on an Agilent Technologies Stratagene Mx3005P qPCR thermocycler (Agilent Technologies, Santa Clara, CA, USA). For each reaction, we used the PCR recipes shown in Table S2 with 1 µl of SYBR Green/ROX solution (one part SYBR Green I nucleic acid gel stain (S7563) (Invitrogen, Carlsbad, CA, USA), four parts ROX Reference Dye (12223-012) (Invitrogen, Carlsbad, CA, USA) and 2000 parts high grade DMSO). Amplifications were carried out with primer specific parameters (Table S6), with 40 cycles followed by a dissociation segment of 95 °C for 1 minute, 52 °C for 30 seconds and 95 °C for 30 seconds. The resulting amplification plots indicated the best dilution factor and volume of template DNA for each primer, as well as the most appropriate number of PCR cycles. In addition, amplification and dissociation curves confirmed that only primer-dimers, and not prey DNA, were present in the extraction blanks.

#### 2.3 PCR amplification

We amplified the targeted sequences using two primer sets, one amplifying a segment of the COI barcode region (Zeale primer set), and the other amplifying a region of the 16S rRNA gene (Epp primer set) (Fig. S2). Both primers were 5' nucleotide tagged (4) to yield a set of unique forward and 60 unique reverse primers. Tags where 7-8 nucleotides in length and there were 2-3 nucleotide mismatches between tags. Each PCR amplification were carried out with matching tags (e.g. F1-R1, F2-R2, etc.) to ensure tag jumps would not result in false assignments of sequences to samples (Schnell et al. 2015). The three PCR replicates from each sample were carried out with different tag combinations to minimise the possible effect of tag bias (5). Each PCR round contained 96 samples, including 90 bat dropping samples, four extraction blanks and two PCR blanks. All PCR mixes were set up in a dedicated pre-PCR laboratory to minimize the risk of contamination. PCR set-up was the same as for qPCR, see above, although omitting the SYBR Green/ROX. PCR reaction recipes and thermal cycler programs are detailed in Table S2 and Table S3 respectively.

#### 2.4 Amplicon visualization, pooling and purification

PCR products were visualised on 2% agarose gels using 4 µl PCR product and always the same loading buffer and visualisation conditions. All blanks appeared negative. For each primer set, multiple amplicon pools were created, which ensured that only PCR products with different tags were pooled to enable sequencing of many PCR replicates in parallel, while being able to track the tagged PCR products back to the correct PCR replicate. Each PCR reaction was assigned an amplification score based on gel band strengths: 3 = very bright band, 2 = bright band, 1 = faint band and 0 = no bad, and these scores were used as a guide for pooling of PCR products: Score 3 = 2 µl, Score 2 = 6 µl, Score 1 = 10 µl, Score 0 = 12 µl. Despite absence of bands in the electrophoresis gels, 10 µl of PCR replicates of blanks were also included. Amplicon pools were subsequently bead-purified using streptavidin coated baits (Beckman Coulter, Brea, CA, USA) and 1:2 amplicon-bead ratio to get rid of primer dimers and eluted in 30 µl of ddH2O.

#### 2.5 Library preparation and sequencing

Amplicon pools were converted into Illumina sequencing libraries using a single-tube library preparation method (6) followed by an indexing PCR to add different reverse indices with at least three nucleotide differences to each amplicon library. The number of indexing PCR cycles was decided based on qPCR screening. Indexed libraries were bead-purified using streptavidin coated baits (Beckman Coulter, Brea, CA, USA) and 1:1.8 amplicon-bead ratio and eluted in 20 µl ddH2O. Purified libraries were analysed in an Agilent Bioanalyzer and combined into sequencing pools using equimolar ratios. Library pools were spiked with 15% PhiX before sequencing them in an Illumina MiSeq platform using 250PE chemistry and aiming 35,000 reads per PCR replicate per sample.

### 3. Metabarcoding bioinformatic procedures

Bioinformatics analyses were carried out in the Danish National Supercomputer for Life Sciences, Computerome. Libraries were demultiplexed in the MiSeq platform after providing the adapter index sequences. The paired end reads in each sequencing library were merged and quality filtered using AdapterRemoval 2.1.7(7). Marker size for Zeale was 158 nucleotides, and for Epp 105 nucleotides. The reads within each library were sorted according to primer and tag sequences using DAMe (8). Samples were subsequently filtered according to the number of replicates in which each sequence was present using DAMe using a relaxed restrictive approach *sensu* Alberdi et al. (9), i.e. only retaining the sequences appearing in at least two of the three PCR replicated. Using a custom script sequences that appeared in the extraction and library blanks of the corresponding batch were removed. Sequences were clustered in OTU with 98% identity, following Alberdi et al. (9), using Sumaclust (10). Samples with less than 5000 sequences were removed, and OTUs with a representation below %0.02 in each sample were removed for their probability of being false positives derived from PCR and sequencing errors. Rarefaction curves and curvature indexes of all samples were generated using the R package DivE 1.0 and samples that neither reached the rarefaction plateau, nor showed a curvature index below 0.80, were discarded. The number of sequences generated per sample was normalized to 0-1 scale using the TSS method. Taxonomy was assigned by aligning the OTU representative sequences to the Genbank nt (11) —and in the case of Zeale also BOLD (12)— databases using Blast+ 2.5.0 suite (13). Finally, the results of both databases were merged using a custom script and selecting the most reliable assignment when finding inconsistencies.

Bayesian OTU phylogenetic trees were generated using BEAST2 (14) after aligning the OTU representative sequences using CLUSTAL Omega (15). All the analyses were performed with a minimal Markov chain Monte Carlo (MCMC) chain length of $10^8$ iterations, sampling trees every 1000. Each Bayesian run was repeated, and convergence of the MCMC chains and sample size was checked using TRACER 1.6.0. To account for the phylogenetic uncertainty of the reconstructed trees, 50 trees were randomly selected from the last 5% (5000) of the trees sampled across the MCMC, as detailed by Alberdi et al. (16).

### 4. Species distribution modelling

Ensemble SDMs were generated for the seven studied bat species using the R package biomod2 (17), including four models (MaxEnt 3.4, Generalized Boosting Model, Random Forests, Flexible Discriminant Analysis). Models included between 113 and 591 occurrence records (Table S5). Species occurrence records were gathered from the online databases GBIF (www.gbif.org) and EUROBATS (https://www.eurobats.org/), from 33 journal publications (Table S6), and unpublished records held by co-authors. To reduce spatial bias and spatial auto-correlation, we first removed low quality records in terms of spatial resolution and taxonomic identification, and then we used the ArcGIS toolbox "SDMtools" (18) to thin spatially clustered records. Models extent was set as the whole of Europe until longitude 42° E to include all the sampling cave locations. Model resolution was set at 30 arc-seconds (~1 km), corresponding to the resolution of the climatic variables.

We initially considered 36 environmental variables (16 climatic, six geographic, 13 habitat and three human disturbance) to include in the models. We tested for correlation among variables, and selected among highly correlated ones ($|r| > 0.75$) the more ecologically relevant or the variable with the stronger effect on model performance on its own for each bat species. Finally, we discarded variables that did not contribute to model performance. Final set of variables used in the models is shown in Tables S1-S2.

Models were run with 10,000 random background points and 1000 maximum iterations. To assess model performance we used tenfold cross-validations replicates, with 75% of records retained for training and 25% for model testing. We used area under the curve (AUC) of the receiver operator characteristics and True Skill Statistic (TSS) to evaluate the models (Table S5 for model performance). The 10 cross-validated replicates were combined to obtain a final predicted environmental suitability map for each of the four modelling methods. Ensemble models were obtained by using AUC values to proportionally weight each method according to its predictive power, excluding models with AUC <0.75.

### 5. Statistical analyses and visualisations

Trophic niche breadth measures and estimations based on Hill numbers were carried out using the R package hilldiv (19). Tests for association between paired samples were performed using the function cor.test based on Pearson's product moment correlation coefficient. The annotated OTU phylogenetic trees were generated with GraPhlan (20), while the rest of the charts were generated with the R package ggplot2. Full codes of the performed analyses are provided in Supplementary Code 3.

### Tables

#### Table S1. **Sampling sites**.

*Miniopterus schreibersii* (MSc), *Myotis capaccinii* (MCa), *Myotis daubentonii* (MDa), *Myotis emarginatus* (MEm), *Myotis myotis* (MMy), *Rhinolophus euryale* (REu) and *Rhinolophus ferrumequinum* (RFe). The resolution of the geographic location data has been lowered in purpose for conservation reasons.

| Site | Country | Latitude | Longitude | Species |
|---|---|---:|---:|---|
| Agua | Spain | 37.32 | -2.16 | MCa |
| Altopiano | Italy | 45.59 | 10.35 | MEm |
| Avenc | Spain | 39.66 | -0.41 | MSc, MCa, Mmy |
| Benevento | Italy | 41.28 | 14.49 | MSc, REu |
| Benimaquia | Spain | 38.82 | 0.06 | MSc, MCa, Mmy |
| Betfia | Romania | 46.98 | 22.02 | MSc |
| Bolera | Spain | 37.73 | -2.92 | RFe |
| Box | United Kingdom | 51.42 | -2.24 | MDa, RFe |
| Campanario | Spain | 37.54 | -6.83 | MDa |
| Clot | Spain | 41.90 | 0.44 | MSc, MCa, Mmy, REu |
| Drenajicka | Serbia | 44.24 | 19.66 | RFe |
| Droves | United Kingdom | 50.93 | -0.76 | MDa |
| Gargina | Bulgaria | 41.85 | 24.93 | MSc, MCa |
| Gesal | Spain | 42.74 | -2.43 | MMy |
| Guixas | Spain | 42.69 | -0.53 | MSc, MEm, REu, RFe |
| Hadzi-Prodanova | Serbia | 43.63 | 20.24 | RFe |
| Heathrow | United Kingdom | 51.49 | -0.51 | MDa |
| Isabel | Spain | 43.26 | -3.37 | MEm, RFe |
| Jasovska | Slovakia | 48.68 | 20.98 | REu, RFe |
| Krokodilos | Greece | 41.22 | 24.72 | MCa |
| Lezate | Spain | 43.30 | -2.55 | MEm, RFe |
| Liliecilor | Romania | 44.47 | 28.48 | MDa |
| Liptsy | Ukraine | 49.54 | 36.43 | MDa |
| Marelli | Italy | 45.86 | 8.78 | MEm |
| Masson | United Kingdom | 53.13 | -1.57 | MDa |
| Montemor | Portugal | 40.14 | -8.71 | MSc, MDa, MEm, RFe |

| | | | | |
|---|---|---|---|---|
| Moura | Portugal | 38.04 | -7.30 | MSc, MMy |
| Ogorelicka | Serbia | 43.35 | 22.09 | MSc, MMy, REu |
| Orlova | Bulgaria | 43.59 | 25.96 | MCa, REu |
| Petnica | Serbia | 44.25 | 19.94 | MSc, MDa, MCa, MMy, REu, RFe |
| Picote | Spain | 37.62 | -6.83 | MDa |
| Reginell | Spain | 42.06 | 3.11 | MEm, RFe |
| Seso | Spain | 42.46 | 0.04 | MDa, REu |
| Soterranya | Spain | 39.68 | -0.49 | MEm, RFe |
| Teglega | Ukraine | 50.20 | 36.37 | MDa |
| Topla | Croatia | 44.19 | 15.84 | MSc, MCa, MMy, REu |
| Toplik | Serbia | 43.76 | 22.31 | RFe |
| Vilar | Spain | 43.07 | -6.95 | MEm |
| Vrasna | Greece | 40.70 | 23.65 | REu |
| Woodchester | United Kingdom | 51.71 | -2.30 | RFe |

#### Table S2. **PCR reaction recipes**.

The concentrations shown in the first column are stock concentrations. The concentrations shown in each of the primer columns are final concentrations.

| **Reagent** | **Zeale** | | **Epp** | |
|---|---|---|---|---|
| dd$H_2O$ | 10.8 µl | | 12.8 µl | |
| Buffer 10x | 2.5 µl | | 2.5 µl | |
| $MgCl_2$ (25 mM) | 2.5 µl | 2.5 mM | 2.5 µl | 2.5 mM |
| dNTP (10 mM each) | 0.5 µl | 0.2 mM | 0.5 µl | 0.2 mM |
| BSA (20 mg/ml) | 1.5 µl | | 1 µl | |
| AmpliTaq Gold® DNA Polymerase (5 U/µl) | 0.2 µl | 1 U | 0.2 µl | 1 U |
| **Total mastermix** | **17** µl | | **19.5** µl | |
| Primer mix (10 µM each) | 4 µl | 1.6 µM | 2.5 µl | 1 µM |
| DNA (1:5 dilution) | 3 µl | | 3 µl | |
| **Total mix** | **25** µl | | **25** µl | |

#### Table S3. **Thermal cycler programs**

| **Zeale** | **Epp** |
|---|---|
| **1x** | **1x** |

| | |
|---|---|
| 95 ºC 10 min | 95 ºC 10 min |
| **40x**<br>95 ºC 20 sec<br>55 ºC 30 sec<br>72 ºC 1 min | **37x**<br>95 ºC 20 sec<br>55 ºC 30 sec<br>72 ºC 30 sec |
| **1x**<br>72 ºC 7 min | **1x**<br>72 ºC 7 min |
| **1x**<br>4 ºC hold | **1x**<br>4 ºC hold |

Table S4. **DNA sequence processing statistics**

| **Statistic** | **Zeale** | **Epp** |
|---|---|---|
| Total sequencing depth | 50.6M | 33.1M |
| Sequencing depth per replicate | 76,356,147±58,211 | 63,041±70,795 |
| Sample size after filtering | 355 | 271 |
| Number of OTUs | 3114 | 1728 |
| Taxonomic (Order) annotation rate | 100% | 100% |
| **Sample sizes after filtering** | **Zeale** | **Epp** |
| ***Miniopterus schreibersii*** | 58 | 46 |
| ***Myotis daubentonii*** | 50 | 43 |
| ***Myotis capaccinii*** | 54 | 44 |
| ***Myotis emarginatus*** | 36 | 21 |
| ***Myotis myotis*** | 41 | 37 |
| ***Rhinolophus euryale*** | 45 | 38 |
| ***Rhinolophus ferrumequinum*** | 63 | 42 |

Table S5. **Species distribution model statistics.**

Number of presence occurrence records (N), variables included in the models (see Table S6 for variable description) and species distribution model performance based on True Skill Statistics (TSS) and Area Under the receiver operator Curve (AUC).

| **Species** | **MSc** | **MDa** | **MCa** | **MEm** | **MMy** | **RFe** | **REu** |
|---|---|---|---|---|---|---|---|
| **N** | **389** | **464** | **113** | **378** | **568** | **591** | **264** |
| **TSS** | **0.78** | **0.64** | **0.87** | **0.74** | **0.71** | **0.72** | **0.81** |
| **AUC** | **0.95** | **0.91** | **0.98** | **0.94** | **0.93** | **0.93** | **0.95** |

| | | | | | | | |
|---|---|---|---|---|---|---|---|
| **Alt** | X | | | X | X | X | X |
| **BIO7** | X | X | | X | X | | X |
| **BIO10** | X | X | X | | X | | X |
| **BIO11** | X | X | X | X | X | X | X |
| **Land** | X | X | X | X | X | X | X |
| **Tree** | X | X | X | X | X | X | X |
| **Het** | | | X | | X | | |
| **River** | X | X | | | X | | |
| **Forest** | X | | X | X | | X | |
| **Grass** | | | X | | | X | |
| **Urban** | | | X | X | | X | |
| **Light** | | X | | | X | | |
| **Karst** | X | | X | X | | X | X |
| **Litho** | X | X | X | X | X | X | X |

Table S6. **Description of the variables used in the species distribution models and their sources.**

| **Type** | **Variable code** | **Variables** | **Source** |
|---|---|---|---|
| | BIO7 | Temperature annual range | www.worldclim.org |
| | BIO10 | Mean temperature of warmest quarter | www.worldclim.org |
| | BIO11 | Mean temperature of coldest quarter | www.worldclim.org |
| Geographic | Alt | Elevation | www.worldclim.org |
| | Karst | Distance to karst | http://arcweb.forest.usf.edu/flex/KarstRegions/ |
| | Litho | Lithology | http://ccgm.org/en/home/168-lithological-map-of-the-world-9782917310250.html |
| Habitat | River | Distance to permanent rivers | www.diva-gis.org |
| | Land | Land cover type | https://earthdata.nasa.gov/ |
| | Forest | Distance to forest | From “Land” |
| | Grass | Distance to grasslands | From “Land ” |
| | Urban | Distance to cities | From “Land” |

| | | | |
|---|---|---|---|
| | Tree | Percentage tree canopy cover (2000) | http://earthenginepartners.appspot.com/science-2013-global-forest/download_v1.1.html |
| | Het | Habitat heterogeneity | From “ Land ” |
| Human impact | Light | Light developing index | http://ngdc.noaa.gov/eog/download.html |

Table S7. **List of publications used to extract species occurrence records.**

| | |
|---|---|
| 1 | Hanak, V., Benda, P., Ruedi, M., Horáček, I. & Sofianidou, T. S. Bats (Mammalia: Chiroptera) of the Eastern Mediterranean, Part 2. New records and review of distribution of bats in Greece. Acta Soc. Zool. Bohemicae 65, 279–346 (2001). |
| 2 | Benda, P. et al. Bats (Mammalia: Chiroptera) of the eastern Mediterranean. Part 3. Review of bat distribution in Bulgaria. Acta Soc. Zool. … 67, 245–357 (2003). |
| 3 | Benda, P. et al. Bats (Mammalia: Chiroptera) of the Eastern Mediterranean. Part Bat fauna of Syria: distribution, systematics, ecology. Acta Soc. Zool. Bohemoslov. / Bohemicae 70, 1–329 (2006). |
| 4 | Benda, P., Hanak, V. & Cerveny, J. Bats (Mammalia: Chiroptera) of the Eastern Mediterranean and Middle East. Part 9. Bats from Transcaucasia and West Turkestan in collection of the National Museum, Prague. Acta Soc. Zool. Bohemicae 75, 159–222 (2011). |
| 5 | Bogdanowicz, W. et al. Cryptic diversity of Italian bats and the role of the Apennine refugium in the phylogeography of the western Palaearctic. Zool. J. Linn. Soc. 174, 635–648 (2015). |
| 6 | Budinski, I., Karapandža, B., Josipović, V., Jovanović, J. & Paunović, M. The first record of alpine long-eared bat Plecotus macrobullaris in Serbia. Turkish J. Zool. 40, 984–988 (2016). |
| 7 | Burazerovic, J. et al. Ticks (Acari: Argasidae, Ixodidae) parasitizing bats in the central Balkans. Exp. Appl. Acarol. 66, 281–291 (2015). |
| 8 | Hamidović, D. Međunarodno Važna Podzemna Skloništa Za Šišmiše Hrvatskoj. (2008). doi:10.13140/RG.2.2.23396.99203 |
| 9 | Hasanspahić, M. & Presetnik, P. Observation of Geoffroy’s bat (Myotis emarginatus) in village Dugo Selo by Olovo (central Bosnia and Herzegovina) during year 2014. Naš krš XXXIV, 29–34 (2014). |
| 10 | Hodžić, M. First finding of the long-fingered bat (Myotis capaccinii) in the cave Bakuf in the village Studenci near Ljubuški. Naš krš XXXV, 12–16 (2015). |
| 11 | Karapandža, B. Details of the first bat (Chiroptera, Mammalia) survey of Vitorog mountain area and the first records of Myotis oxygnathus monticelli, 1885, Myotis Bechsteinii (kuhl, 1817) and Barbastella barbastellus (Schreber, 1774) in Bosnia and Herzegovina. Naš krš XXXIV, 3–10 (2014). |
| 12 | Branko Karapandža. Details of the first bat (Chiroptera, Mammalia) survey of Mišarica cave near Banjaluka town. Hypsugo I, 12–19 (2016). |
| 13 | Kipson, M. Strengthening the support and scientific evidence for conservation of " Europe ’ s Amazon " through monitoring of bats as bioindicators and involvement of community. (2012). |
| 14 | Mazija, M. & Rnjak, D. Survey results of selected bat roost sites in Popovo Polje within the Ravno municipality (Bosnia and Herzegovina). Hypsugo I, 20–29 (2016). |
| 15 | Nikola, M., Prestenik, P., Branko, M. & Martin, C. Contribution to the knowledge of the Macedonian |

| | |
|---|---|
| | bat fauna. Vespertilio 17, 103–114 (2014). |
| 16 | Mulaomerović, J. & Dervović, T. Two Mediterranean bat species from cave Peruc at village Izbišno (SE B&H). Naš krš XXXV, 25–26 (2015). |
| 17 | Nagy, Z. L. & Postawa, T. Seasonal and geographical distribution of cave-dwelling bats in Romania: Implications for conservation. Anim. Conserv. 14, 74–86 (2011). |
| 18 | Pašić, J. & Presetnik, P. Daubenton' s bat (Myotis daubentonii (Kuhl, 1817)) new species on the list of bats (Chiroptera) of Bosnia and Herzegovina. Naš Krš XXXIII, 8–13 (2013). |
| 19 | Pašić, J. & Presetnik, P. Second record of Daubenton' s bat (Myotis daubentonii (Kuhl, 1817)) and second and further records of Kuhl's pipistrelle (Pipistrellus kuhlii, (Kuhl, 1817)) in Bosnia and Hercegovina. Naš krš XXXIV, 11–15 (2014). |
| 20 | Jasmin Pašić, Jasminko Mulaomerović, P. P. Results of survey of potential bat hibernacula in Bosnia and Herzegovina in winter 2012/13. Naš Krš XXXIII, 23–34 (2013). |
| 21 | Pavlinić, I., Đaković, M. & Tvrtković, N. The Atlas of Croation Bats (Chiroptera) Part I. Nat. Croat. 19, 295–337 (2010). |
| 22 | Primož, P. et al. Distribution of bats (Chiroptera) in Montenegro. Vespertilio 17, 129–156 (2014). |
| 23 | Presetnik, P., Mulaomerović, J. & Pašić, J. Results of survey of potential bat hibernacula in Bosnia and Herzegovina in winter 2014/15. Hypsugo 1, 30–37 (2016). |
| 24 | Presetnik, P. Results of bat and other mammals fauna survey on VI. Internacionalni Biology camp "Stolac 2016" (Bosnia i Herzegovina). Hypsugo II, 17–26 (2017). |
| 25 | Presetnik, P. et al. Survey results of potential bat hibernacula in Bosnia and Herzegovina in winter 2016/17. Hypsugo II, 27–41 (2017). |
| 26 | Presetnik, P., Radonjić, M., Pavlovič, E., Gojznikar, J. & Jovanović, M. Results of bat survey during biology students research camp "Ekosistemi Balkana – Skadarsko Jezero 2017" (Montenegro). Hypsugo II, 41–52 (2017). |
| 27 | Rnjak, D., Rnjak, G., Hanžek, N. & Zrnčić1, V. Bat fauna research at the foot of Velež mountain (Bosnia and Herzegovina) in 2014. Hypsugo II, 11–30 (2017). |
| 28 | Rnjak, D., Goran, R. & Zrnčić, V. Bat fauna research in Šibenik, Unešić and Drniš municipalities, 2013 – 2014. Hypsugo I, 9–24 (2016). |
| 29 | Sachanowicz, K., Ciechanowski, M., Rachwald, A. & Piskorski, M. Overview of bat species reported in Albania with the first country records for eight species. J. Nat. Hist. 2933, 9p. (2015). |
| 30 | Théou, P., Loce, E. & Đurović, M. Results of the pioneer survey of potential bat hibernacula in Albania (2012 – 2015). Nat. Slov. 17, 25–39 (2015). |
| 31 | Uhrin, M., Benda, P., Obuch, J. & Urban, P. Changes in abundance of hibernating bats in central Slovakia (1992–2009). Biologia (Bratisl). 65, 349–361 (2010). |
| 32 | Uhrin, M. et al. Revision of the occurrence of Rhinolophus euryale in the Carpathian region, Central Europe. Vespertilio 16, 289–328 (2012). |

##### Table S8. **Primer-specific relation between trophic (dRER) and spatial niche breadth**

| **Primer** | **Metric** | **Pearsons' r** | **T statistic** | **df** | **p-value** |
|---|---|---|---|---|---|
| **Zeale** | **B1** | 0.85 | 30.154 | 348 | < 0.001 |

| | | | | | |
|---|---|---|---|---|---|
| | **B2** | 0.78 | 23.081 | 348 | < 0.001 |
| **Epp** | **B1** | 0.82 | 26.987 | 348 | < 0.001 |
| | **B2** | 0.89 | 36.566 | 348 | < 0.001 |

Table S9. **Relation between behavioral traits and spatial niche breadth**

| Trait | Metric | Pearsons' r | T statistic | df | p-value |
|---|---|---|---|---|---|
| **Hunting** | **B1** | 0.59 | 1.655 | 5 | 0.1588 |
| | **B2** | 0.58 | 1.6025 | 5 | 0.17 |
| **Habitat** | **B1** | -0.70 | 2.1687 | 5 | 0.08227 |
| | **B2** | -0.57 | -1.5396 | 5 | 0.1843 |
| **Roosting** | **B1** | -0.48 | -1.2332 | 5 | 0.2723 |
| | **B2** | -0.40 | -0.99054 | 5 | 0.3674 |

Table S10. **Relation between behavioral traits and dietary niche breadth**

| Trait | Metric | Pearsons' r | T statistic | df | p-value |
|---|---|---|---|---|---|
| **Hunting** | **dR** | -0.53 | -1.4097 | 5 | 0.2177 |
| | **dRE** | 0.44 | 1.0911 | 5 | 0.325 |
| | **dRER** | 0.79 | 24.347 | 348 | < 0.001 |
| **Habitat** | **dR** | 0.48 | 1.231 | 5 | 0.2731 |
| | **dRE** | -0.38 | -0.93139 | 5 | 0.3944 |
| | **dRER** | -0.58 | -13.302 | 348 | < 0.001 |
| **Roosting** | **dR** | -0.24 | -0.56439 | 5 | 0.5969 |
| | **dRE** | -0.33 | -0.79223 | 5 | 0.4641 |
| | **dRER** | -0.08 | -1.6391 | 348 | 0.1021 |

Table S11. **Correlation between observed and estimated diversities**

| Primer | Measure | F-statistic | DF | $r^2$ | p-value |
|---|---|---|---|---|---|
| Zeale | dR | 129 | 1, 19 | 0.86 | <0.001 |
| Zeale | dRER | 931.7 | 1, 19 | 0.98 | <0.001 |
| Epp | dR | 62.64 | 1, 19 | 0.76 | <0.001 |
| Epp | dRER | 201.8 | 1, 19 | 0.91 | <0.001 |

Table S12. **Hunting strategy traits**

Hunting strategies: A = aerial hawking, T = trawling, G = gleaning, F = flycatching

| Species | A | T | G | F | Reference |
|---|---|---|---|---|---|
| ***Miniopterus schreibersii*** | 10 | 0 | 0 | 0 | Presetnik & Aulagnier 2013 (21) |
| ***Miniopterus schreibersii*** | 10 | 0 | 0 | 0 | Vincent et al. 2010 (22) |
| ***Miniopterus schreibersii*** | 10 | 0 | 0 | 0 | Norberg & Rayner 1987 (23) |
| ***Myotis daubentonii*** | 8 | 2 | 0 | 0 | Todd & Waters 2007 (24) |
| ***Myotis daubentonii*** | 5 | 5 | 0 | 0 | Geberl et al. 2015 (25) |
| ***Myotis daubentonii*** | 5 | 5 | 0 | 0 | Kalko and Schnitzler 1989 (26) |
| ***Myotis capaccinii*** | 5 | 5 | 0 | 0 | Ahlen & Rydell 1990 (27) |
| ***Myotis capaccinii*** | 0 | 9 | 1 | 0 | Siemers 2001 (28) |
| ***Myotis capaccinii*** | 1 | 9 | 0 | 0 | Biscardi 2007 (29) |
| ***Myotis emarginatus*** | 5 | 0 | 5 | 0 | Krull et al. 1991 (30) |
| ***Myotis emarginatus*** | 8 | 0 | 2 | 0 | Goiti et al. 2011 (31) |
| ***Myotis emarginatus*** | 0 | 0 | 10 | 0 | Dekker et al. 2013 (32) |
| ***Myotis emarginatus*** | 3 | 0 | 7 | 0 | Schumm et al. 1991 (33) |
| ***Myotis myotis*** | 0 | 0 | 9 | 1 | Norberg and Rayner 1987 (23) |
| ***Myotis myotis*** | 3 | 0 | 7 | 0 | Arlettaz 1996 (34) |
| ***Myotis myotis*** | 2 | 0 | 8 | 0 | Audet 1990 (35) |
| ***Myotis myotis*** | 1 | 0 | 8 | 1 | Fenton 1990 (36) |
| ***Rhinolophus euryale*** | 8 | 0 | 0 | 2 | Goiti et al. 2003 (37) |
| ***Rhinolophus euryale*** | 10 | 0 | 0 | 0 | Siemers & Ivanova 2004 (38) |
| ***Rhinolophus euryale*** | 9 | 0 | 0 | 1 | Russo et al. 2002 (39) |
| ***Rhinolophus ferrumequinum*** | 5 | 0 | 0 | 5 | Jin et al. 2005 (40) |
| ***Rhinolophus ferrumequinum*** | 7 | 0 | 0 | 3 | Jones & Rayner 1989 (41) |
| **AVERAGE** | | | | | |
| **Species** | **A** | **T** | **G** | **F** | |
| ***Miniopterus schreibersii*** | 10 | 0 | 0 | 0 | |
| ***Myotis daubentonii*** | 6 | 4 | 0 | 0 | |
| ***Myotis capaccinii*** | 2 | 7.67 | 0.33 | 0 | |
| ***Myotis emarginatus*** | 4 | 0 | 6 | 0 | |
| ***Myotis myotis*** | 1.5 | 0 | 8 | 0.5 | |
| ***Rhinolophus euryale*** | 9 | 0 | 0 | 1 | |

| | | | | |
|---|---|---|---|---|
| ***Rhinolophus ferrumequinum*** | 6 | 0 | 0 | 4 |

Table S13. **Habitat use traits**

Habitat types: O = open (e.g. meadows, pastures, arable land, bareland), S = semi-open (e.g. open forest, orchards), F = forest (e.g. broadleaf, coniferous), W = water (e.g. rivers, streams, ponds), U = urban.

| **Species** | **O** | **S** | **F** | **W** | **U** | Reference |
|---|---|---|---|---|---|---|
| ***Miniopterus schreibersii*** | 2 | 1 | 1 | 1 | 5 | Vincent et al. 2010 (22) |
| ***Myotis daubentonii*** | 0 | 0 | 0 | 10 | 0 | Siivonen & Wermundsen 2008 (42) |
| ***Myotis daubentonii*** | 0 | 0 | 2 | 8 | 0 | Ahlen & Rydell 1990 (27) |
| ***Myotis daubentonii*** | 0 | 0 | 0 | 10 | 0 | Dietz et al. 2006 (43) |
| ***Myotis capaccinii*** | 0 | 0 | 0 | 10 | 0 | Almenar et al. 2009 (44) |
| ***Myotis capaccinii*** | 0 | 0 | 0 | 10 | 0 | Biscardi et al. 2007 (29) |
| ***Myotis emarginatus*** | 0 | 4 | 6 | 0 | 0 | Flaquer et al. 2008 (45) |
| ***Myotis emarginatus*** | 0 | 2 | 7 | 0 | 1 | Krull et al. 1991 (30) |
| ***Myotis emarginatus*** | 0 | 2 | 8 | 0 | 0 | Goiti et al. 2011 (31) |
| ***Myotis myotis*** | 4 | 3 | 3 | 0 | 0 | Arlettaz 1999 (46) |
| ***Myotis myotis*** | 1 | 1 | 8 | 0 | 0 | Audet 1990 (35) |
| ***Myotis myotis*** | 1 | 3 | 6 | 0 | 0 | Zahn et al. 2005 (47) |
| ***Myotis myotis*** | 1 | 5 | 3 | 0 | 1 | Drescher 2004 (48) |
| ***Myotis myotis*** | 0 | 1 | 9 | 0 | 0 | Rudolph et al. 2009 (49) |
| ***Rhinolophus euryale*** | 0 | 1 | 9 | 0 | 0 | Goiti et al. 2003 (37) |
| ***Rhinolophus euryale*** | 1 | 3 | 6 | 0 | 0 | Russo et al. 2002 (39) |
| ***Rhinolophus euryale*** | 0 | 2 | 8 | 0 | 0 | Russo et al. 2005 (50) |
| ***Rhinolophus ferrumequinum*** | 2 | 3 | 5 | 0 | 0 | Flanders & Jones 2009 (51) |
| **AVERAGES** | | | | | | |
| **Species** | **O** | **S** | **F** | **W** | **U** | |
| ***Myotis daubentonii*** | 0 | 0 | 0.67 | 9.33 | 0 | |
| ***Myotis capaccinii*** | 0 | 0 | 0 | 10 | 0 | |
| ***Myotis emarginatus*** | 0 | 2.67 | 7 | 0 | 0.33 | |
| ***Myotis myotis*** | 1.4 | 2.6 | 5.8 | 0 | 0.2 | |
| ***Miniopterus schreibersii*** | 2 | 1 | 1 | 1 | 5 | |
| ***Rhinolophus euryale*** | 0.33 | 2 | 7.67 | 0 | 0 | |
| ***Rhinolophus ferrumequinum*** | 2 | 3 | 5 | 0 | 0 | |

Table S14. **Roosting traits**

Roost types: U = underground cavities, B = buildings, T = trees, C = crevices

| Species | U | B | T | C | Reference |
|---|---|---|---|---|---|
| ***Miniopterus schreibersii*** | 8 | 2 | 0 | 0 | Benda & Paunović 2019 (52) |
| ***Myotis daubentonii*** | 0 | 2 | 5 | 2 | Bogdanowicz 1994 (53) |
| ***Myotis daubentonii*** | 0 | 0 | 10 | 0 | Boonman 2000 (54) |
| ***Myotis daubentonii*** | 0 | 0 | 10 | 0 | Encarnaçao et al. 2005 (55) |
| ***Myotis daubentonii*** | 0 | 0 | 10 | 0 | Kapfer et al. 2007 (56) |
| ***Myotis capaccinii*** | 10 | 0 | 0 | 0 | Papadatou et al. 2009 (57) |
| ***Myotis capaccinii*** | 9 | 1 | 0 | 0 | Almenar 2006 (58) |
| ***Myotis emarginatus*** | 0 | 9 | 1 | 0 | Krull et al. 1991 (30) |
| ***Myotis emarginatus*** | 7 | 3 | 0 | 0 | Karataş & Özgül 2003 (59) |
| ***Myotis emarginatus*** | 5 | 5 | 0 | 0 | Zahn et al. 2010 (60) |
| ***Myotis myotis*** | 7 | 2 | 0 | 1 | Zahn 1999 (61) |
| ***Rhinolophus euryale*** | 9 | 1 | 0 | 0 | Budinski et al. 2019 (62) |
| ***Rhinolophus euryale*** | 8 | 2 | 0 | 0 | Uhrin et al. 2012 (63) |
| ***Rhinolophus ferrumequinum*** | 6 | 4 | 0 | 0 | Dietz et al. 2013 (64) |
| **AVERAGES** | | | | | |
| **Species** | **U** | **B** | **T** | **C** | |
| ***Miniopterus schreibersii*** | 8 | 2 | 0 | 0 | |
| ***Myotis daubentonii*** | 0 | 0.5 | 8.75 | 0.5 | |
| ***Myotis capaccinii*** | 9.5 | 0.5 | 0 | 0 | |
| ***Myotis emarginatus*** | 4 | 5.67 | 0.33 | 0 | |
| ***Myotis myotis*** | 7 | 2 | 0 | 1 | |
| ***Rhinolophus euryale*** | 8.5 | 1.5 | 0 | 0 | |
| ***Rhinolophus ferrumequinum*** | 6 | 4 | 0 | 0 | |

### Figures

Figure S1. **Trophic spectrum of the analysed species as retrieved from the Zeale primers**

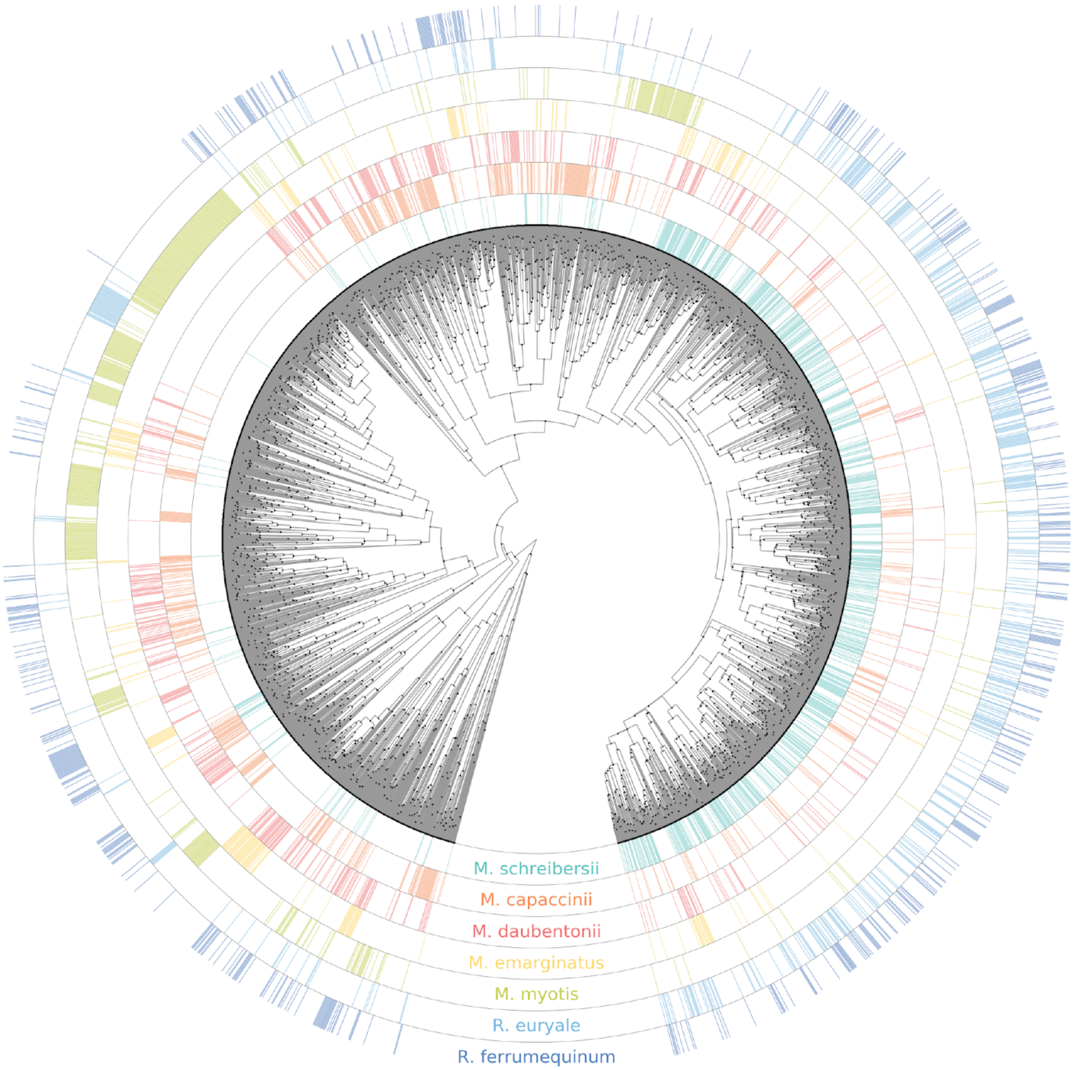

Figure S2. **Trophic spectrum of the analysed species as retrieved from the Epp primers**

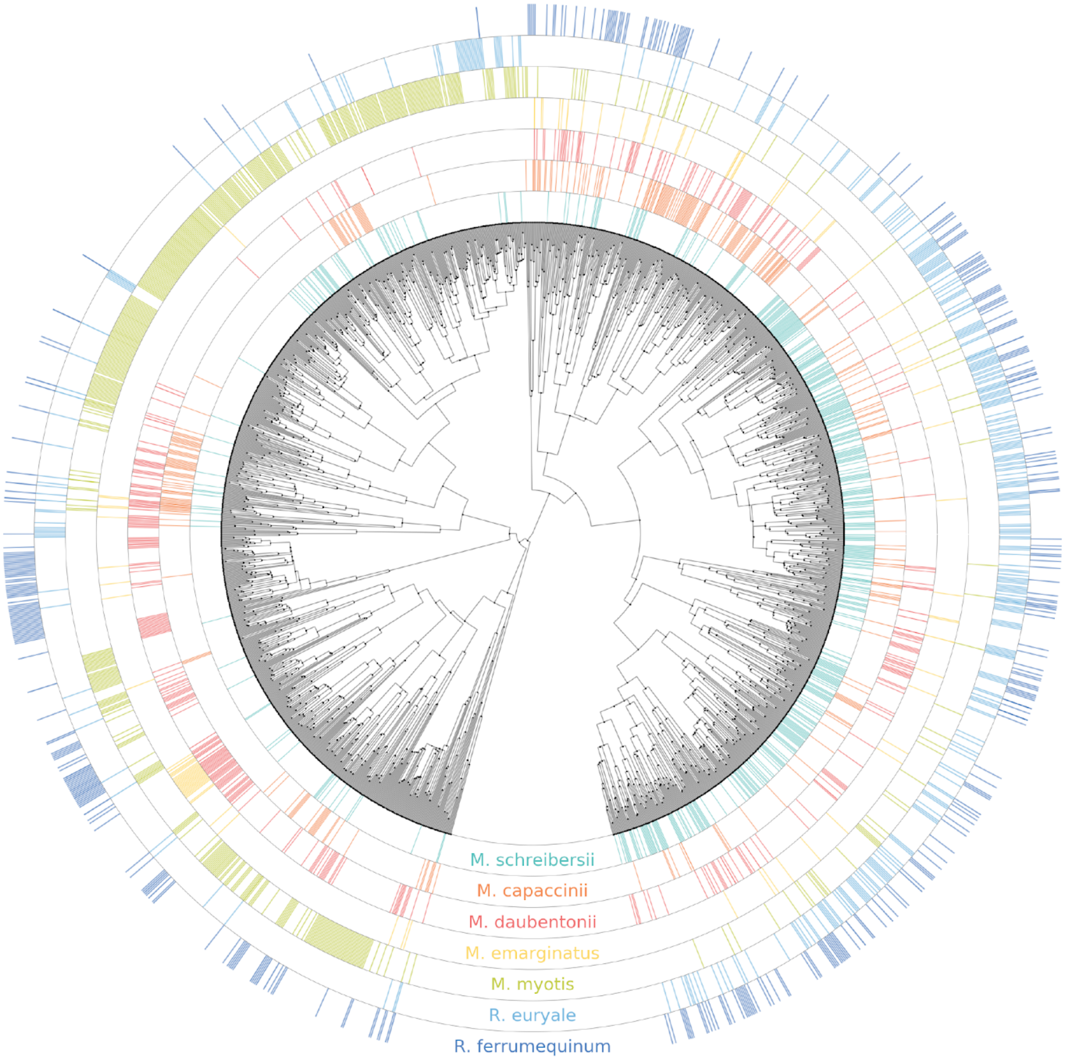

Figure S3. **Spatial projections of the niche models**

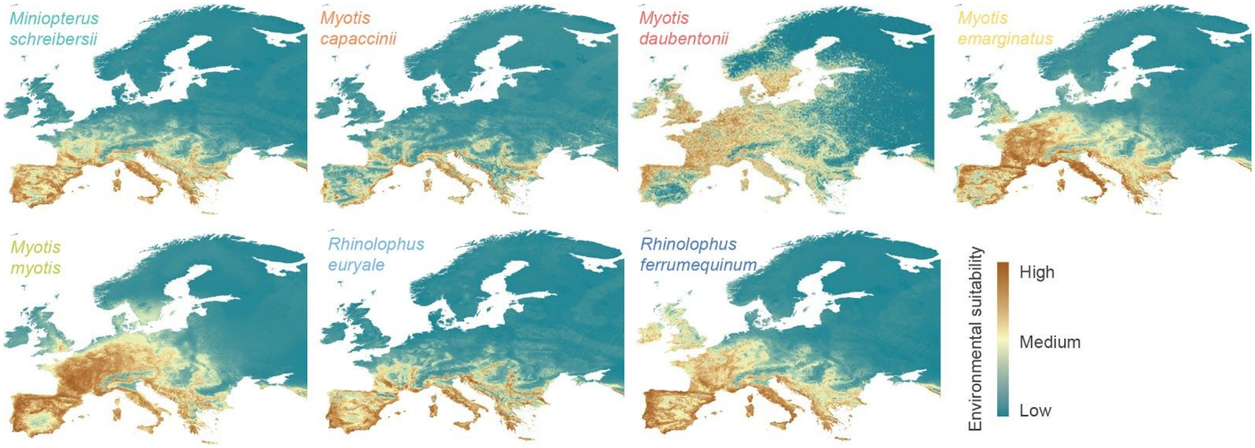

Figure S4. **Relation of species trophic niche breadth measures with spatial niche breadth**

A) Trophic niche breadth measures derived from the Zeale (A) and Epp (B) datasets.

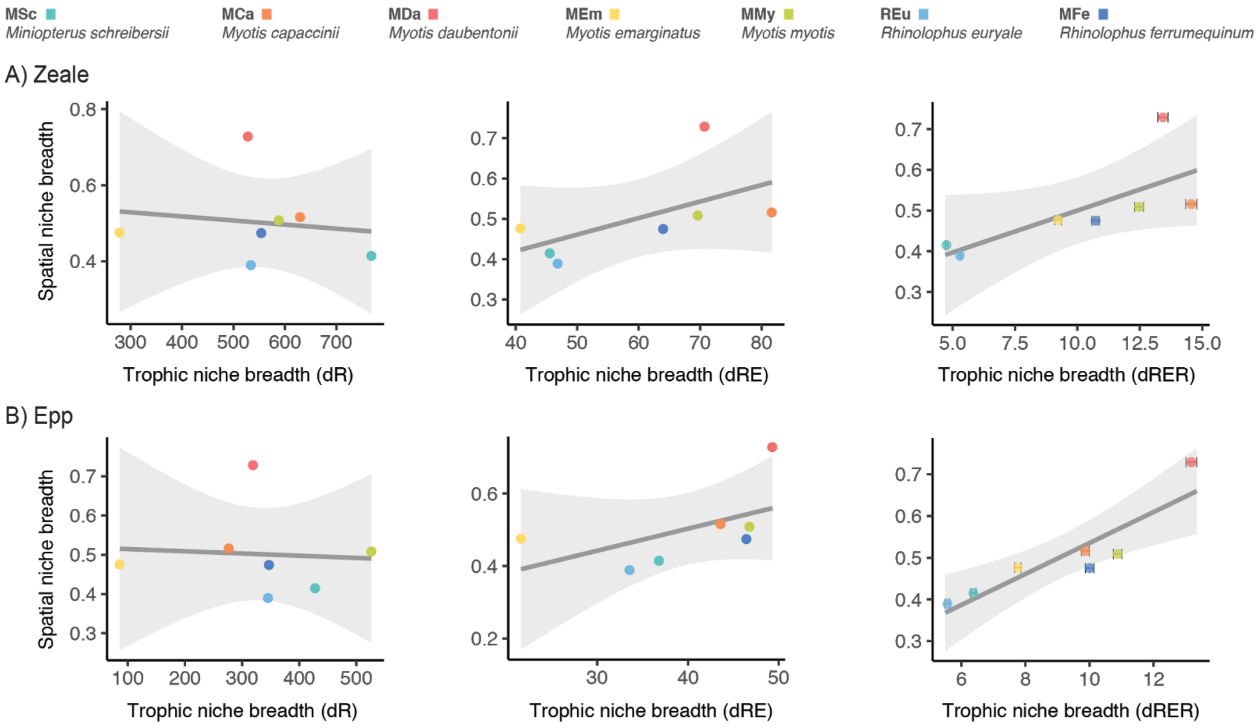
